# Marine bacterial enrichment in the sea surface microlayer, and surface taxa aerosolization potential in the Western Mediterranean Sea

**DOI:** 10.1101/2023.04.26.538450

**Authors:** Julie Dinasquet, Birthe Zäncker, Alessia Nicosia, Estelle Bigeard, Anne-Claire Baudoux, Anja Engel, Cecile Guieu, Ingrid Obernosterer, Karine Sellegri

## Abstract

The sea surface microlayer (SSML) is critical to air-sea exchanges of gases and primary aerosols. However, despite the extent of this boundary layer, little is known about its specific bacterial community (bacterioneuston) and how it may affect ocean-atmosphere exchanges. Here, we studied the bacterial community composition in the surface waters of three different basins of the Western Mediterranean Sea and assessed the selective air-sea transfer of marine bacteria through experimental nascent sea spray aerosol production in a 10 L tank with plunging jets. In situ, the bacterioneuston harbored basin-specific enriched taxa and followed a similar spatial pattern as the underlying bacterioplankton community. Aerosolization potential showed that sea spray taxa might be recruited from both the underlying water and the SSML, and that taxa enriched in the bacterioneuston were not always aerosolized. Our results suggest that the Mediterranean nutrient gradient, as well as pulse events such as dust deposition, affect the distribution of the bacterial community at the ocean-atmosphere interface, which may impact biogeochemical processes, climate regulation and bacterial dispersal through aerosolization.

## Introduction

The sea surface microlayer (SSML) is a thin (1-1000 µm) gel-like matrix at the surface of the ocean (Engel *et al*., 2017 and reference therein). This ocean skin harbors a specific ecology and chemistry in comparison to the underlying water (ULW). The biogeochemical and physical processes there, are critical to air-sea exchanges of energy, gases and primary aerosols which affect climate (Engel *et al*., 2017, Barthelmeß *et al*., 2021, Hendrickson *et al*., 2021).

Atmospheric deposition can also accumulate at the SSML, influencing biogeochemical processes (Astrahan *et al*., 2016) before being exported to deeper waters. However, despite the geographic extent of the SSML, the biogeographic distribution of the bacterial community living there (the bacterioneuston) and its potential role in ocean-atmosphere connectivity is still poorly understood.

The bacterioneuston is a key player in the SSML biogeochemical cycles and in air-sea processes. Its activities affect organic matter composition and concentration, gas fluxes, physical properties of the SSML through gel formation, and thus, aerosol composition and numbers (e.g. Upstill-Goddard *et al*., 2003, Reinthaler *et al*., 2008, Kurata *et al*., 2016, Engel *et al*., 2018, Freney *et al*., 2020, Sun *et al*., 2020, Sellegri *et al*., 2021). The bacterioneuston composition appears to be tightly linked to the ULW bacterial community. Nevertheless, while some studies found that the bacterioneuston was enriched in specific taxa compared to the ULW (Franklin *et al*., 2005, Zäncker *et al*., 2018), others report no significant differences between the two layers (Agogue *et al*., 2005, Obernosterer *et al*., 2008). These discrepancies might be the results of spatiotemporal variations, winds, and SSML chemical properties but also be the results of the SSML collection method (Stolle *et al*., 2010, Cunliffe *et al*., 2011, Rahlff *et al*., 2017, Wong *et al*., 2021).

Sea spray aerosols (SSA) represents one of the most abundant sources of natural aerosol particles in the atmosphere and contribute significantly to global aerosol fluxes (McNeill, 2017). SSA are produced mainly by rising bubbles bursting at the air-sea interface. As bubbles rise through the water column they scavenge inorganic and biological material, which play a significant role in the SSML formation (Engel *et al*., 2017). Thus, bacterial accumulation at the SSML might be a precursor to their aerosolization; and SSML properties might affect the selective transfer of bacteria to SSA (Aller *et al*., 2005, Michaud *et al*., 2018). Indeed, nascent SSA have been shown to contain a large diversity of bacteria which appear to be a subset of the community present in the surface water (Fahlgren *et al*., 2015, Rastelli *et al*., 2017, Michaud *et al*., 2018, Freitas *et al*., 2022). These studies suggest that bacterial aerosolization is selective, which might be influenced by cellular properties and environmental conditions (Rastelli *et al*., 2017, Michaud *et al*., 2018). Some of these airborne bacteria can directly influence atmospheric processes, cloud formation and climate through their activities and icenucleating capacities (Amato *et al*., 2017, Failor *et al*., 2017, Beall *et al*., 2021, Trueblood *et al*., 2021, Šantl-Temkiv *et al*., 2022). Moreover, as SSA travel long distance (Bondy *et al*., 2017, Mayol *et al*., 2017, Šantl-Temkiv *et al*., 2018, Johansson *et al*., 2019), they are a potential vector of microbial dispersal, influencing bacterial biogeography.

Here, we studied the bacterial community composition in the surface waters of three different basins of the oligotrophic Western Mediterranean Sea. We compared the spatial distribution of the bacterioneuston with the bacterioplankton in the ULW, and evaluated the taxa enriched in the SSML between the studied basins. We further assessed the aerosolization potential of surface taxa through onboard experimental nascent SSA production.

## Material and Methods

### Sea water sampling

Seawater was collected during the ‘ProcEss studies at the Air-sEa Interface after dust deposition in the MEditerranean sea’ cruise (PEACETIME), onboard the R/V “Pourquoi Pas ?” in May/June 2017 (Guieu *et al*., 2020). Stations were sampled in the Ionian Sea (ION), Tyrrhenian Sea (TYRR), and in the Algerian basin (WEST) from a zodiac (Fig. S1). Water collection is described in detail in Zäncker *et al*. (2021). Briefly, at each station sea surface microlayer (SSML) water was sampled using the glass plate method, and uUnderlying water (ULW) was sampled 20 cm below the SSML with two acid-cleaned bottles that were opened ca. 20 cm below the surface. At station FAST (the western most station), we sampled surface water before a wet dust deposition event, and then 2h, 24h and 48h after the rain (SSML was not sampled before the rain).

### Nascent sea spray aerosols

Nascent sea spray aerosols (nSSA) were produced onboard, as described in Trueblood *et al*. (2021). Briefly, nSSA were generated in a 10 L glass tank with a plunging jet system linked to a continuous flow of surface water (5 m) from the ship underway system. nSSA filters were collected for bacterial community composition at two stations in the Ionian Sea, and three stations in the Algerian basin (Fig. S1),. and at station FAST 2h and 24h after the wet dust deposition event. nSSA were filtered onto glass fiber filters over 24h at 16.5 LPM and immediately stored at -20°C. Samples for bacterial abundance in nSSA were collected from 28 stations using a PILS sampler. 0.5 mL samples were fixed with glutaraldehyde (0.5% final), flash frozen and stored at -80°C.

### Bacterial abundance

2mL samples were fixed with glutaraldehyde (1% final concentration) for bacterial abundance in the SSML and the ULW. Samples were stored immediately at -20°C. Abundance was measured by flow-cytometry as described in Zäncker *et al*. (2021).

For bacterial abundance in the nSSA, the samples were stained with Sybr Green and measured by flow cytometry (BD-FACS Calibur). True count beads and 2µm beads were used as reference. Potential autofluorescent particles were subtracted from the counts.

### DNA sampling, sequencing and sequence analysis

400 mL water for SSML and ULW were filtered through a 0.2µm filter (Durapore) after a pre-filtration through a 100 µm mesh, and immediately stored at -80°C. DNA was extracted as described in Zäncker *et al*. (2021). DNA extracts were quantified and normalized and used as templates for PCR amplification of the V4-V5 region of the 16S rRNA gene (Parada *et al*., 2016), as described in Dinasquet *et al*. (2022). The nSSA filters were extracted with a Zymo quick Fungal/Bacterial DNA miniprep as described in Dadaglio *et al*. (2018). DNA extracts were quantified and normalized and used as templates for PCR amplification of the V4 region of the 16S rRNA gene (Apprill *et al*., 2015).

All reads were processed using the Quantitative Insight Into Microbial Ecology 2 pipeline (QIIME2 v2020.2, Bolyen *et al*., 2019). Reads were truncated to 250bp to cover the V4 region, based on sequencing quality, denoised, merged and chimera-checked using DADA2 (Callahan *et al*., 2016). A total of 953 amplicon sequence variants (ASVs) were obtained. Taxonomy assignments were made against the database SILVA 132 (Quast *et al*., 2013). All sequences associated with this study have been deposited under the BioProject ID: PRJNA693966.

### Statistics

Alpha and beta-diversity indices for community composition were estimated after randomized subsampling to 7888 reads for the analysis with nSSA samples and to 24728 for the analysis with only the surface water samples16S rDNA. Analyses were run in QIIME 2 and in Primer v.6 software package (Clarke & Warwick, 2001). Differences between the samples richness and diversity were assessed using Kruskal-Wallis pairwise test. Beta diversities were run on Bray Curtis dissimilarity. Differences between samples’ beta diversity were tested using PERMANOVA (Permutational Multivariate Analysis of Variance) with pairwise test and 999 permutations. The sequences contributing most to the dissimilarity between layers were identified using the ANalysis of Composition of Microbiomes (ANCOM)(Mandal *et al*., 2015).

## Results and discussion

The Mediterranean Sea is the largest semi-enclosed ocean basin. It is also a large Low-Nutrient-Low-Chlorophyll (LNLC) region, characterized by a long summer stratification period and a west-to-east gradient of increasing oligotrophy, where microbial growth is generally limited by phosphorus availability (Durrieu de Madron *et al*., 2011). The Mediterranean bacterial biogeography was shown to reflect the oligotrophy gradient and physical barriers from surface to deep waters (Mapelli *et al*., 2013, Sebastián *et al*., 2021). Here, we further describe, for the first time to our knowledge, the spatial distribution of the Mediterranean bacterial community at the air-sea interface and the aerosolization potential of surface taxa.

### Spatial distribution of bacteria community composition in the SSML and ULW

The biogeochemical context of the SSML sampled during this cruise is described in Tovar-Sanchez et al. (2020) and Zäncker et al. (2021). Briefly, these studies show that trace metal and TEP were consistently enriched in the SSML. DOC was also enriched in the SSML (as shown here, Fig.S2). Picophytoplankton were generally more abundant in the SSML, but no significant difference was observed in the eukaryotic community composition and diversity (Zäncker et al, 2021). The bacterial abundance decreased from west to east in both the SSML and ULW but showed no significant enrichment in the SSML (Table 1, Tovar-Sánchez *et al*., 2020, Zäncker *et al*., 2021).The bacterioplankton community, however, showed significant differences between layers. Evenness was significantly lower in the SSML compared to the ULW (Fig.S3, Kruskal Wallis p=0.006).

**Table 1:**
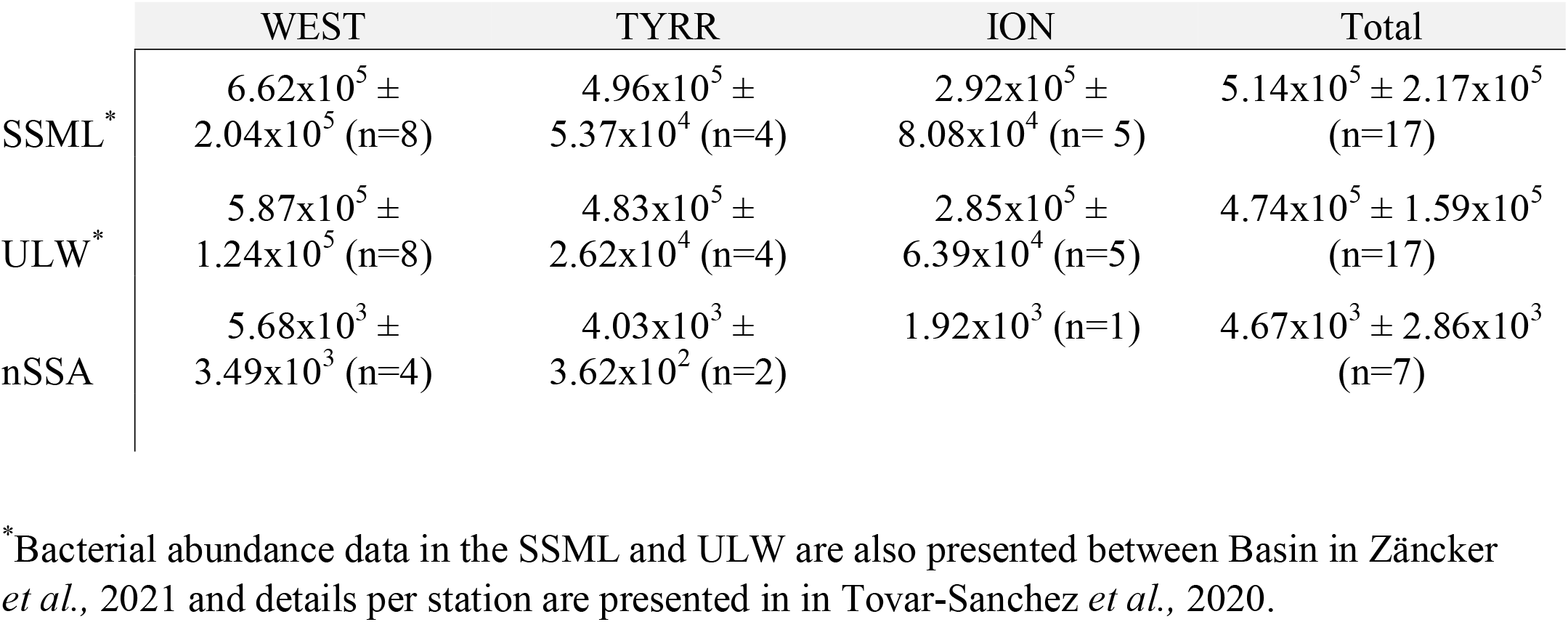
Bacterial abundance in the three sampled basins, in the nascent sea spray aerosols (cells m^-3^) and in the surface water (SSML and ULW, cells mL^-1^).

The bacterioneuston has been shown to be tightly linked to the bacterial community in the ULW just below (Cunliffe *et al*., 2013). Overall, reflecting previous studies from surface mixed layer communities, the spatial distribution of the bacterial community in the first micrometers of the SSML followed the Mediterranean Sea oligotrophic gradient and physical barriers. The overall surface bacterioplankton (combined SSML and ULW) was significantly different between basins (Fig.1, Permanova p<0.01). Similar spatial distribution of the main taxa reported for surface mixed layers (Mapelli *et al*., 2013, Sebastián *et al*., 2021) were also observed here.

**Figure 1:**
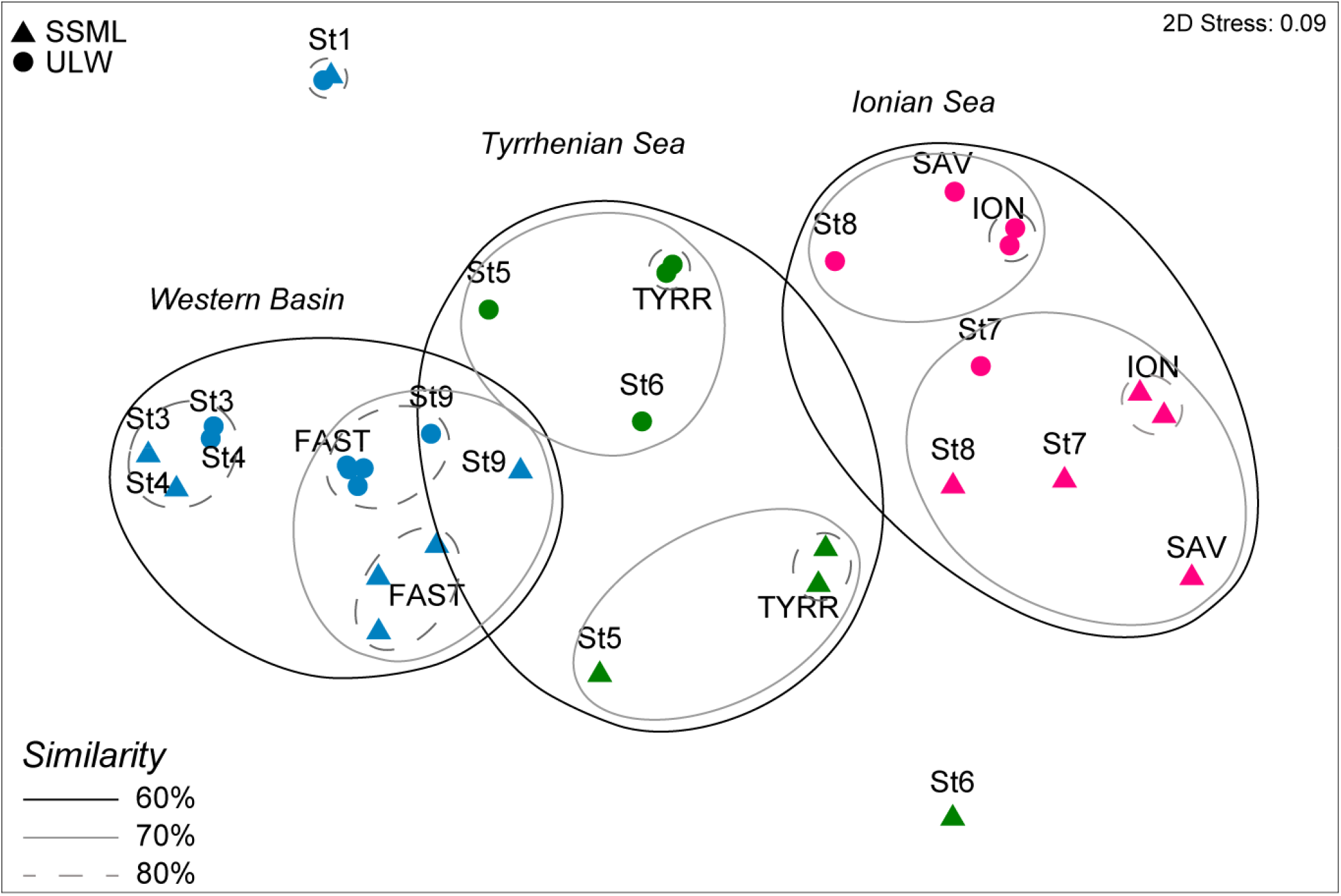
nMDS plot of bacterial community composition in the SSML and ULW at each station based on Bray-Curtis dissimilarities of the 16S rDNA sequences. Samples clustering at different level of similarity are circled together, basin groups are significantly different from each other (p < 0.01) based on a PERMANOVA test.

Indeed, the relative abundance of the Gamma-Proteobacteria, Oceanospirillales, Alteromonadales and Pseudoalteromonadales increased from West-to-East in both the SSML and ULW, while the relative abundance of cyanobacteria (*Synechococcus*, but especially *Prochlorococcus*) decreased (Fig.2, S4).

**Figure 2:**
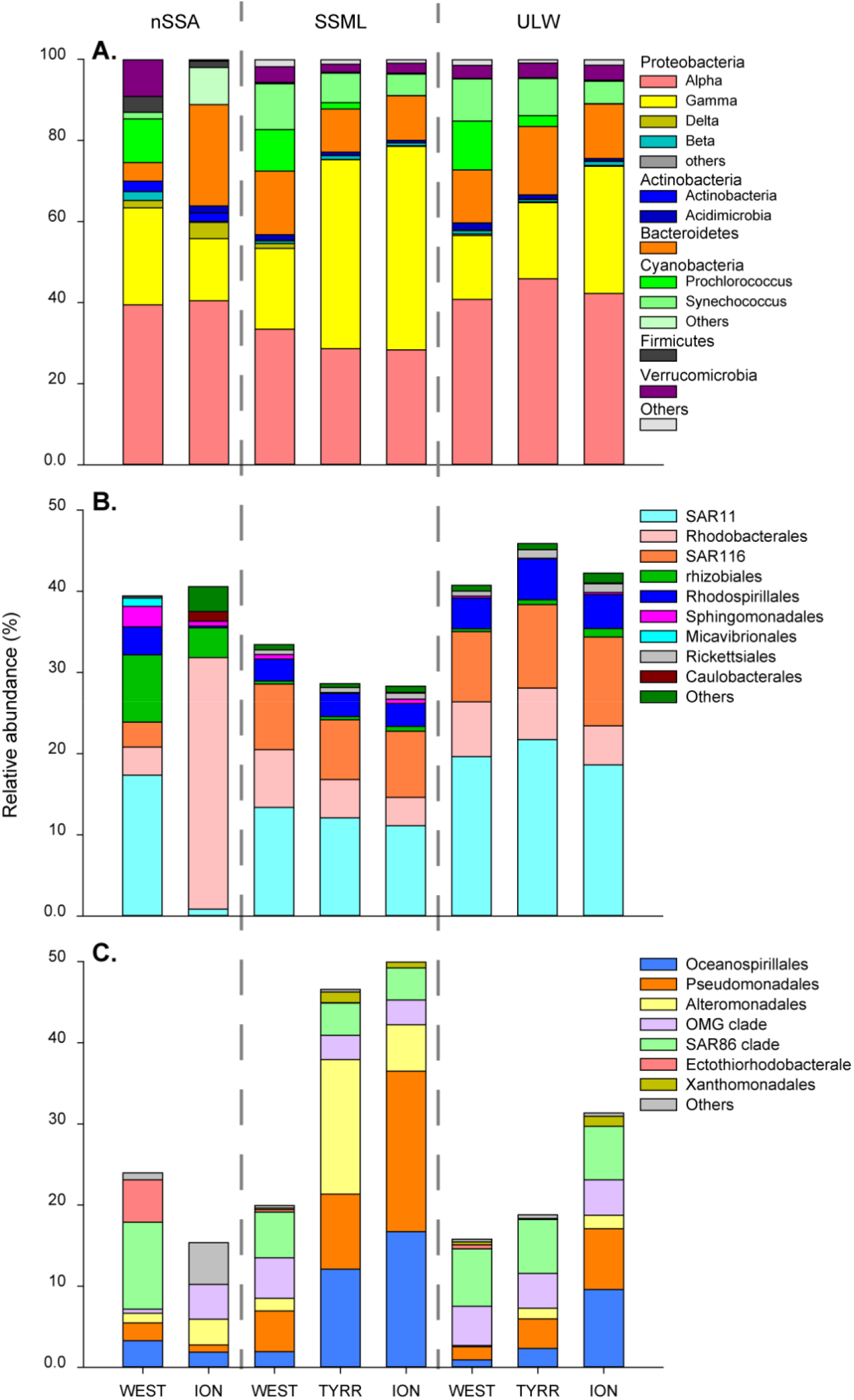
Average community composition at the phylum level (A), Alpha-proteobacteria family level (B) and Gamma-proteobacteria family level (C) distribution in the different basins in the nSSA, SML and ULW.

Nevertheless, due to the weak cloud coverage, the Mediterranean bacterioneuston is subject to strong solar radiation that may influence its composition compared to lower surface layers.

The bacterioneuston has only been studied at a few coastal stations of the North-west Mediterranean Sea (Agogue *et al*., 2005, Joux *et al*., 2006), and despite strong potential stressors no significant differences were observed between the ULW and the SSML. Here, closely comparing the ULW and SSML communities showed that the bacterioneuston in the Ionian and Tyrrhenian Seas were significantly different between the two layers (Fig.1, p<0.05). Generalist Gamma-Proteobacteria (Oceanospirillales, Alteromonadales and Pseudoalteromonadales) were particularly enriched in the SSML compared to the ULW (Fig.2, S4), which may be due to different physico-chemical properties of the SSML.

No significant differences were observed between SSML and ULW communities in the Western basin. However, at station FAST (Fig.S1) which was influenced by a wet dust deposition event (Tovar-Sánchez *et al*., 2020, Van Wambeke *et al*., 2020, Desboeufs *et al*., 2022), the SSML and ULW clustered separately with specific taxa changing in response to dust compared to other Western stations (Fig.1, S4). This might be due to the Mediterranean bacteria’s capacity to rapidly respond to dust input (Tsiola *et al*., 2017, Dinasquet *et al*., 2022), especially in the SSML (Astrahan *et al*., 2016) or to the direct deposition of dust attached bacteria (Rahav *et al*., 2016, Mescioglu *et al*., 2019, Aalismail *et al*., 2020) to the SSML.

### Aerosolization potential of surface taxa to sea spray aerosols

It is challenging to disentangle the role of marine bacteria on the atmospheric microbiome due to the large influence of bacteria attached to anthropogenic, terrestrial and mineral aerosols (e.g.Lang-Yona *et al*., 2022), in particular in the semi-enclosed and dust impacted Mediterranean Sea (Rahav *et al*., 2016, Gat *et al*., 2017, Mescioglu *et al*., 2019). To isolate the aerosolization potential of marine bacteria, complex experimental procedures with nSSA production are necessary. But only few studies have investigated the selective transfer of marine taxa from surface waters into nSSA (Fahlgren *et al*., 2015, Rastelli *et al*., 2017, Michaud *et al*., 2018, Freitas *et al*., 2022).

Sea spray aerosols are formed by bubbles rising through the water column and potentially scavenging material from the surface and SSML. Hence, bacterial accumulation at the SSML might be a precursor to their aerosolization (Aller *et al*., 2005).In order to better understand the aerosolization potential of bacteria in the surface Mediterranean Sea, we compared the sequential enrichment of bacterial taxa from the ULW to the SSML, to onboard experimentally produced nSSA with surface water from the same stations. Concentration of bacteria in the nSSA likely followed the same west-to-east gradient of decreasing abundance as the ULW and SSML (Table 1) and were in line with previous observations in marine aerosols in situ, in the Mediterranean Sea (from 7 to 25×103 cells m-3)(Mescioglu *et al*., 2019) and in the North Atlantic (∼8.02×10^3^ cells m^-3^)(Mayol *et al*., 2014). The nSSA bacterial community showed significantly lower diversity but higher evenness than the surface communities (Fig. S3, Kruskal-Wallis p <0.001). The community structure also appeared to be basin specific with the Ionian and Western nSSA communities clustered separately from ULW and SSML water communities (Fig. S5), suggesting selective transfers of bacterial taxa, as observed in previous studies (Fahlgren *et al*., 2015, Rastelli *et al*., 2017, Michaud *et al*., 2018, Freitas *et al*., 2022). Interestingly the transferred cyanobacteria followed a similar spatial gradient as in surface layers, with *Prochlorococcus* better aerosolized in the Western basin where it is more abundant and *Synechococcus* in the Ionian Sea. Sequential enrichment of some taxa (e.g. Rhizobiales, *Sphingobium, Pseudomonas, Vibrio* and *Staphylococcus*) was observed from ULW, SSML to nSSA but generally aerosolized taxa appeared to be present but not abundant throughout the surface waters. Some taxa that were enriched in the SSML had a low aerosolization potential, which suggests that not all bacterioneuston taxa are scavenged into the aerosols, and this might be specific to cellular properties or environmental factors especially the physical properties of the SSML.

Most of the aerosolized taxa observed here (Table 2, Fig 2, 3, S4) are commonly observed in marine aerosols in other oceanic regions (e.g. Cho & Hwang, 2011, Fahlgren *et al*., 2015, Michaud *et al*., 2018, Mescioglu *et al*., 2019, Lang-Yona *et al*., 2022), suggesting that cellular properties consistent within these groups independent from environmental factors, influence their sea-to-air transfer. A few of these taxa (Table 2, Fig 2, 3, S4) are related to known opportunistic pathogens (e.g. *Vibrio, Pseudomonas, Staphylococcus, Streptococcus*) for marine organisms.

**Table 2:** Enrichment in the SSML and aerosolization potential in the nSSA and of the most abundant genus responsible for the difference between the communities in the ULW, SSML and nSSA based on Analysis of composition of microbiome (ANCOM).

| Class | Order | Genus | SSML<br>enrichment | Aerosolization<br>potential |  |
| --- | --- | --- | --- | --- | --- |
|  |  |  | From ULW | from<br>SSML | from<br>ULW |
| Alpha-proteobacteria | Pelagibacterales | SAR11 clade | 52% | 189% | 98% |
| Alpha-proteobacteria | Puniceispirillales | SAR116 clade | 108% | 31% | 33% |
| Alpha-proteobacteria | Rhizobiales | *Bradyrhizobium* | 111% | 53300% | 59012% |
| Alpha-proteobacteria | Rhizobiales | *Methylobacterium* | 260% | 1662% | 4320% |
| Alpha-proteobacteria | Rhodobacterales | *Ascidiahabitans* | 61% | 47% | 29% |
| Alpha-proteobacteria | Rhodospirillales | AEGEAN-169 | 60% | 182% | 110% |
| Alpha-proteobacteria | Sphingomonadales | *Sphingobium* | 720% | 639% | 4598% |
| Beta-proteobacteria | Burkholderiales | *Delftia* | 63% | 1924% | 1216% |
| Delta-proteobacteria | Bdellovibrionales | OM27 clade | 448% | 91% | 407% |
| Gamma-proteobacteria | Alteromonadales | *Pseudoalteromonas* | 542% | 12% | 67% |
| Gamma-proteobacteria | Alteromonadales | *Alteromonas* | 246% | 17% | 41% |
| Gamma-proteobacteria | Cellvibrionales | OM60 clade | 88% | 18% | 16% |
| Gamma-proteobacteria | Oceanospirillales | *Halomonas* | 363% | 1% | 3% |
| Gamma-proteobacteria | Oceanospirillales | *Litoricola* | 79% | 4% | 3% |
| Gamma-proteobacteria | Oceanospirillales | *Marinomonas* | 81% | 25029% | 20154% |
| Gamma-proteobacteria | Pseudomonadales | *Acinetobacter* | 152% | 9% | 13% |
| Gamma-proteobacteria | Pseudomonadales | *Pseudomonas* | 197% | 181% | 356% |
| Gamma-proteobacteria | Vibrionales | *Vibrio* | 173% | 2739% | 4732% |
| Gamma-proteobacteria | SAR86 clade |  | 72% | 491% | 352% |
| Gamma-proteobacteria | Xanthomonadales | *Stenotrophomonas* | 233% | 0% | 0% |
| Bacilli | Bacillales | *Staphylococcus* | 146% | 8975% | 13140% |
| Bacteroidetes | Balneolales | *Balneola* | 76% | 74% | 56% |
| Bacteroidetes | Flavobacteriales | NS4 | 64% | 23% | 14% |
| Bacteroidetes | Flavobacteriales | NS9 | 92% | 91% | 84% |
| Bacteroidetes | Flavobacteriales | NS5 | 85% | 51% | 43% |
| Cyanophyceae | Synechococcales | *Prochlorococcus* | 71% | 202% | 142% |
| Cyanophyceae | Synechococcales | *Synechococcus* | 77% | 19% | 14% |
| Verrucomicrobia | Opitutales | *Coralimargarita* | 69% | 354% | 252% |
| Verrucomicrobia | Opitutales | *Lentimonas* | 79% | 181% | 147% |

**Figure 3:**
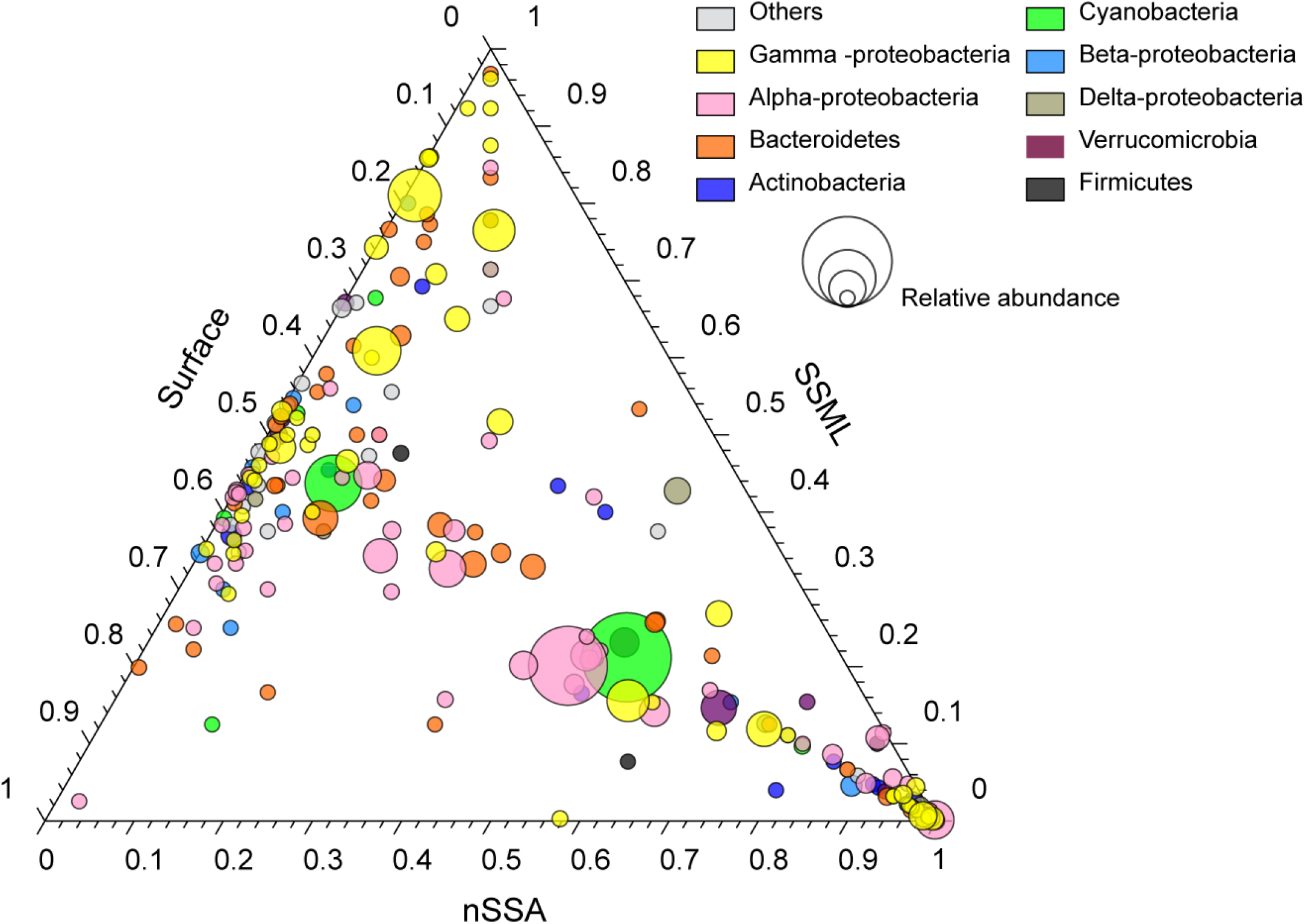
Ternary plot bacterial taxa enriched across the three fractions: surface water, sea surface microlayer (SSML) and nascent sea spray aerosol (nSSA). Each bubble represents one major bacterial group (collapsed at the genus level colored per phylum), the size of the bubble represents the overall relative abundance of the group in the three fractions. “Others” is an average of low abundance phyla.

Others are associated to known phototrophs or have specific sulfur requirements (e.g. SAR11, SAR86, *Ectothiorhodobacterales, Coraliomargarita*), which may be an advantage to survive harsh atmospheric conditions. Some of the taxa with high aerosolization potentials have also been shown to increase in response to higher temperature or to dust deposition (e.g.Westrich *et al*., 2016, Sato-Takabe *et al*., 2019, Dinasquet *et al*., 2022), which suggests that their dispersal may have a greater impact on marine biogeochemical cycles and ecosystem health in the future Mediterranean Sea.

### Conclusion

The biogeographic distribution of the bacterial community in the very first centimeters of the water column – the sea-air interface, is influenced by similar horizontal and vertical gradients, as previously described for lower depths. This surface community appears to follow the oligotrophic gradient but may also be influenced by pulse events like dust deposition that stimulate the growth of specific opportunistic taxa. The bacterioneuston plays a crucial role in the ocean and the atmosphere connectivity, where they are critical to air-sea exchange affecting climate. Here, we show through onboard experiments that surface Mediterranean taxa have a range of aerosolization potential. As described in other oceanic regions, these taxa can be sequentially enriched through the surface layers and selectively transferred into the aerosols.

Sequential enrichment in the SSML and aerosolization of bacteria may be affected by future climate conditions and aerosol deposition in the Mediterranean Sea. This could have a large impact on climate, microbial biogeography and ecosystem services.

## Funding

This work was supported by a Marie Skłodowska-Curie Actions International Postdoctoral Fellowship (grant no. PIOF-GA-2013-629378) to JD. This study is a contribution to the PEACETIME project (http://peacetime-project.org, https://doi.org/10.17600/17000300), a joint initiative of the MERMEX and ChArMEx components supported by CNRS-INSU, IFREMER, CEA, and Météo-France as part of the program MISTRALS, which is coordinated by INSU. PEACETIME was endorsed as a process study by GEOTRACES and SOLAS.

## Acknowledgements

We gratefully acknowledge the onboard support from the captain and crew of the R/V *Pourquoi Pas?* and our chief scientists Cécile Guieu and Karine Desboeuf. We also thank Barbara Marie for the analysis of DOC concentration, and Tania Klüver for flow cytometry analyses.

## Conflict of interest statement

The authors declare that the research was conducted in the absence of any conflict of interest.

## Notes

### Competing Interest Statement

The authors have declared no competing interest.

## References

Aalismail NA, Díaz-Rúa R, Ngugi DK, Cusack M & Duarte CM (2020) Aeolian Prokaryotic Communities of the Global Dust Belt Over the Red Sea. Frontiers in Microbiology 11.

Agogue H, Casamayor EO, Bourrain M, Obernosterer I, Joux F, Herndl GJ & Lebaron P (2005) A survey on bacteria inhabiting the sea surface microlayer of coastal ecosystems. Fems Microbiology Ecology 54: 269–280.

Aller JY, Kuznetsova MR, Jahns CJ & Kemp PF (2005) The sea surface microlayer as a source of viral and bacterial enrichment in marine aerosols. Journal of Aerosol Science 36: 801–812.

Amato P, Joly M, Besaury L, Oudart A, Taib N, Moné AI, Deguillaume L, Delort A-M & Debroas D (2017) Active microorganisms thrive among extremely diverse communities in cloud water. PLOS ONE 12: e0182869.

Apprill A, McNally S, Parsons R & Weber L (2015) Minor revision to V4 region SSU rRNA 806R gene primer greatly increases detection of SAR11 bacterioplankton. Aquatic Microbial Ecology 75: 129–137.

Astrahan P, Herut B, Paytan A & Rahav E (2016) The Impact of Dry Atmospheric Deposition on the Sea-Surface Microlayer in the SE Mediterranean Sea: An Experimental Approach. Frontiers in Marine Science 3.

Barthelmeß T, Schütte F & Engel A (2021) Variability of the Sea Surface Microlayer Across a Filament’s Edge and Potential Influences on Gas Exchange. Frontiers in Marine Science 8.

Beall CM, Michaud JM, Fish MA, Dinasquet J, Cornwell GC, Stokes MD, Burkart MD, Hill TC, DeMott PJ & Prather KA (2021) Cultivable halotolerant ice-nucleating bacteria and fungi in coastal precipitation. Atmospheric Chemistry and Physics 21: 9031–9045.

Bolyen E & Rideout JR & Dillon MR, et al. (2019) Reproducible, interactive, scalable and extensible microbiome data science using QIIME 2. Nature Biotechnology 37: 852–857.

Bondy AL, Wang B, Laskin A, Craig RL, Nhliziyo MV, Bertman SB, Pratt KA, Shepson PB & Ault AP (2017) Inland Sea Spray Aerosol Transport and Incomplete Chloride Depletion: Varying Degrees of Reactive Processing Observed during SOAS. Environmental Science & Technology 51: 9533–9542.

Callahan BJ, McMurdie PJ, Rosen MJ, Han AW, Johnson AJA & Holmes SP (2016) DADA2: Highresolution sample inference from Illumina amplicon data. Nature Methods 13: 581.

Cho BC & Hwang CY (2011) Prokaryotic abundance and 16S rRNA gene sequences detected in marine aerosols on the East Sea (Korea). FEMS Microbiology Ecology 76: 327–341.

Clarke KR & Warwick PE (2001) Change in Marine Communities: An Approach to Statistical Analysis and Interpretation.

Cunliffe M, Upstill-Goddard RC & Murrell JC (2011) Microbiology of aquatic surface microlayers 1. Fems Microbiology Reviews 35: 233–246.

Cunliffe M, Engel A, Frka S, Gasparovic B, Guitart C, Murrell JC, Salter M, Stolle C, Upstill-Goddard R & Wurl O (2013) Sea surface microlayers: A unified physicochemical and biological perspective of the air-ocean interface. Progress in Oceanography 109: 104–116.

Dadaglio L, Dinasquet J, Obernosterer I & Joux F (2018) Differential responses of bacteria to diatom-derived dissolved organic matter in the Arctic Ocean. Aquatic Microbial Ecology 82: 59–72.

Desboeufs K, Fu F, Bressac M, et al. (2022) Wet deposition in the remote western and central Mediterranean as a source of trace metals to surface seawater. Atmos Chem Phys 22: 2309–2332.

Dinasquet J, Bigeard E, Gazeau F, Azam F, Guieu C, Marañón E, Ridame C, Van Wambeke F, Obernosterer I & Baudoux AC (2022) Impact of dust addition on the microbial food web under present and future conditions of pH and temperature. Biogeosciences 19: 1303–1319.

Durrieu de Madron X, Guieu C, Sempéré R, et al. (2011) Marine ecosystems’ responses to climatic and anthropogenic forcings in the Mediterranean. Progress in Oceanography 91: 97–166.

Engel A, Sperling M, Sun C, Grosse J & Friedrichs G (2018) Organic Matter in the Surface Microlayer: Insights From a Wind Wave Channel Experiment. Frontiers in Marine Science 5.

Engel A, Bange HW, Cunliffe M, et al. (2017) The Ocean’s Vital Skin: Toward an Integrated Understanding of the Sea Surface Microlayer. Frontiers in Marine Science 4.

Fahlgren C, Gómez-Consarnau L, Zábori J, Lindh MV, Krejci R, Mårtensson EM, Nilsson D & Pinhassi J (2015) Seawater mesocosm experiments in the Arctic uncover differential transfer of marine bacteria to aerosols. Environmental Microbiology Reports 7: 460–470.

Failor KC, Schmale Iii DG, Vinatzer BA & Monteil CL (2017) Ice nucleation active bacteria in precipitation are genetically diverse and nucleate ice by employing different mechanisms. ISME J.

Franklin MP, McDonald IR, Bourne DG, Owens NJP, Upstill-Goddard RC & Murrell JC (2005) Bacterial diversity in the bacterioneuston (sea surface microlayer): the bacterioneuston through the looking glass. Environmental Microbiology 7: 723–736.

Freitas GP, Stolle C, Kaye PH, Stanley W, Herlemann DPR, Salter ME & Zieger P (2022) Emission of primary bioaerosol particles from Baltic seawater. Environmental Science: Atmospheres 2: 1170–1182.

Freney E, Sellegri K, Nicosia A, et al. (2020) Mediterranean nascent sea spray organic aerosol and relationships with seawater biogeochemistry. Atmos Chem Phys Discuss 2020: 1–23.

Gat D, Mazar Y, Cytryn E & Rudich Y (2017) Origin-Dependent Variations in the Atmospheric Microbiome Community in Eastern Mediterranean Dust Storms. Environmental Science & Technology 51: 6709–6718.

Guieu C, D’Ortenzio F, Dulac F, Taillandier V, Doglioli A, Petrenko A, Barrillon S, Mallet M, Nabat P & Desboeufs K (2020) Introduction: Process studies at the air–sea interface after atmospheric deposition in the Mediterranean Sea – objectives and strategy of the PEACETIME oceanographic campaign (May–June 2017). Biogeosciences 17: 5563–5585.

Hendrickson BN, Brooks SD, Thornton DCO, Moore RH, Crosbie E, Ziemba LD, Carlson CA, Baetge N, Mirrielees JA & Alsante AN (2021) Role of Sea Surface Microlayer Properties in Cloud Formation. Frontiers in Marine Science 7.

Johansson JH, Salter ME, Acosta Navarro JC, Leck C, Nilsson ED & Cousins IT (2019) Global transport of perfluoroalkyl acids via sea spray aerosol. Environmental Science: Processes & Impacts 21: 635–649.

Joux F, Agogué H, Obernosterer I, Dupuy C, Reinthaler T, Herndl G J. & Lebaron P (2006) Microbial community structure in the sea surface microlayer at two contrasting coastal sites in the northwestern Mediterranean Sea. Aquatic Microbial Ecology 42: 91–104.

Kurata N, Vella K, Hamilton B, Shivji M, Soloviev A, Matt S, Tartar A & Perrie W (2016) Surfactant-associated bacteria in the near-surface layer of the ocean. Scientific Reports 6: 19123.

Lang-Yona N, Flores JM, Haviv R, et al. (2022) Terrestrial and marine influence on atmospheric bacterial diversity over the north Atlantic and Pacific Oceans. Communications Earth & Environment 3: 121.

Mandal S, Van Treuren W, White RA, Eggesbø M, Knight R & Peddada SD (2015) Analysis of composition of microbiomes: a novel method for studying microbial composition. Microbial Ecology in Health and Disease 26: 27663.

Mapelli F, Varela MM, Barbato M,Alvariño R, Fusi M, Álvarez M, Merlino G, Daffonchio D & Borin S (2013) Biogeography of planktonic bacterial communities across the whole Mediterranean Sea. Ocean Sci 9: 585–595.

Mayol E, Jimenez MA, Herndl GJ, Duarte CM & Arrieta JM (2014) Resolving the abundance and air-sea fluxes of airborne microorganisms in the North Atlantic Ocean. Frontiers in Microbiology 5.

Mayol E, Arrieta JM, Jiménez MA, et al. (2017) Long-range transport of airborne microbes over the global tropical and subtropical ocean. Nature Communications 8: 201.

McNeill VF (2017) Atmospheric Aerosols: Clouds, Chemistry, and Climate. Annual Review of Chemical and Biomolecular Engineering 8: 427–444.

Mescioglu E, Rahav E, Belkin N, Xian P, Eizenga JM, Vichik A, Herut B & Paytan A (2019) Aerosol Microbiome over the Mediterranean Sea Diversity and Abundance. Atmosphere 10: 440.

Michaud JM, Thompson LR, Kaul D, et al. (2018) Taxon-specific aerosolization of bacteria and viruses in an experimental ocean-atmosphere mesocosm. Nature Communications 9: 2017.

Obernosterer I, Catala P, Lami R, Caparros J, Ras J, Bricaud A, Dupuy C, Van Wambeke F & Lebaron P (2008) Biochemical characteristics and bacterial community structure of the sea surface microlayer in the South Pacific Ocean. Biogeosciences 5: 693–705.

Parada AE, Needham DM & Fuhrman JA (2016) Every base matters: assessing small subunit rRNA primers for marine microbiomes with mock communities, time series and global field samples. Environmental Microbiology 18: 1403–1414.

Quast C, Pruesse E, Yilmaz P, Gerken J, Schweer T, Yarza P, Peplies J & Glöckner FO (2013) The SILVA ribosomal RNA gene database project: improved data processing and web-based tools. Nucleic Acids Research 41: D590–D596.

Rahav E, Ovadia G, Paytan A & Herut B (2016) Contribution of airborne microbes to bacterial production and N2 fixation in seawater upon aerosol deposition. Geophysical Research Letters 43: 719–727.

Rahlff J, Stolle C, Giebel H-A, Brinkhoff T, Ribas-Ribas M, Hodapp D & Wurl O (2017) High wind speeds prevent formation of a distinct bacterioneuston community in the sea-surface microlayer. FEMS Microbiology Ecology 93.

Rastelli E, Corinaldesi C, Dell’Anno A, Lo Martire M, Greco S, Cristina Facchini M, Rinaldi M, O’Dowd C, Ceburnis D & Danovaro R (2017) Transfer of labile organic matter and microbes from the ocean surface to the marine aerosol: an experimental approach. Scientific Reports 7: 11475.

Reinthaler T, Sintes E & Herndl GJ (2008) Dissolved organic matter and bacterial production and respiration in the sea □ surface microlayer of the open Atlantic and the western Mediterranean Sea. Limnology and Oceanography 53: 122–136.

Šantl-Temkiv T, Gosewinkel U, Starnawski P, Lever M & Finster K (2018) Aeolian dispersal of bacteria in southwest Greenland: their sources, abundance, diversity and physiological states. FEMS Microbiology Ecology 94.

Šantl-Temkiv T, Amato P, Casamayor EO, Lee PKH & Pointing SB (2022) Microbial ecology of the atmosphere. FEMS Microbiology Reviews.

Sato-Takabe Y, Hamasaki K & Suzuki S (2019) High temperature accelerates growth of aerobic anoxygenic phototrophic bacteria in seawater. MicrobiologyOpen 8: e00710–e00710.

Sebastián M, Ortega-Retuerta E, Gómez-Consarnau L, Zamanillo M, Álvarez M, Arístegui J & Gasol JM (2021) Environmental gradients and physical barriers drive the basin-wide spatial structuring of Mediterranean Sea and adjacent eastern Atlantic Ocean prokaryotic communities. Limnology and Oceanography n/a.

Sellegri K, Nicosia A, Freney E, et al. (2021) Surface ocean microbiota determine cloud precursors. Scientific Reports 11: 281.

Stolle C, Nagel K, Labrenz M & Jürgens K (2010) Succession of the sea-surface microlayer in the coastal Baltic Sea under natural and experimentally induced low-wind conditions. Biogeosciences 7: 2975–2988.

Sun H, Zhang Y, Tan S, Zheng Y, Zhou S, Ma Q-Y, Yang G-P, Todd JD & Zhang X-H (2020) DMSP-producing bacteria are more abundant in the surface microlayer than subsurface seawater of the East China Sea. Microbial ecology 80: 350–365.

Tovar-Sánchez A, Rodríguez-Romero A, Engel A, et al. (2020) Characterizing the surface microlayer in the Mediterranean Sea: trace metal concentrations and microbial plankton abundance. Biogeosciences 17: 2349–2364.

Trueblood JV, Nicosia A, Engel A, et al. (2021) A two-component parameterization of marine icenucleating particles based on seawater biology and sea spray aerosol measurements in the Mediterranean Sea. Atmos Chem Phys 21: 4659–4676.

Tsiola A, Tsagaraki TM, Giannakourou A, Nikolioudakis N, Yücel N, Herut B & Pitta P (2017) Bacterial Growth and Mortality after Deposition of Saharan Dust and Mixed Aerosols in the Eastern Mediterranean Sea: A Mesocosm Experiment. Frontiers in Marine Science 3.

Upstill-Goddard RC, Frost T, Henry GR, Franklin M, Murrell JC & Owens NJP (2003) Bacterioneuston control of air-water methane exchange determined with a laboratory gas exchange tank. Global Biogeochemical Cycles 17: /a-n/a.

Van Wambeke F, Taillandier V, Deboeufs K, Pulido-Villena E, Dinasquet J, Engel A, Marañón E, Ridame C & Guieu C (2020) Influence of atmospheric deposition on biogeochemical cycles in an oligotrophic ocean system. Biogeosciences Discuss 2020: 1–51.

Westrich JR, Ebling AM, Landing WM, Joyner JL, Kemp KM, Griffin DW & Lipp EK (2016) Saharan dust nutrients promote Vibrio bloom formation in marine surface waters. Proceedings of the National Academy of Sciences 113: 5964–5969.

Wong S-K, Suzuki S, Cui Y, Kaneko R, Kogure K & Hamasaki K (2021) Sampling Constraints and Variability in the Analysis of Bacterial Community Structures in the Sea Surface Microlayer. Frontiers in Marine Science 8.

Zäncker B, Cunliffe M & Engel A (2018) Bacterial Community Composition in the Sea Surface Microlayer Off the Peruvian Coast. Frontiers in Microbiology 9.

Zäncker B, Cunliffe M & Engel A (2021) Eukaryotic community composition in the sea surface microlayer across an east–west transect in the Mediterranean Sea. Biogeosciences 18: 2107–2118.

