## Supplementary material for "Marine bacterial enrichment in the sea surface microlayer, and surface taxa aerosolization potential in the Western Mediterranean Sea": Supplemtary figures

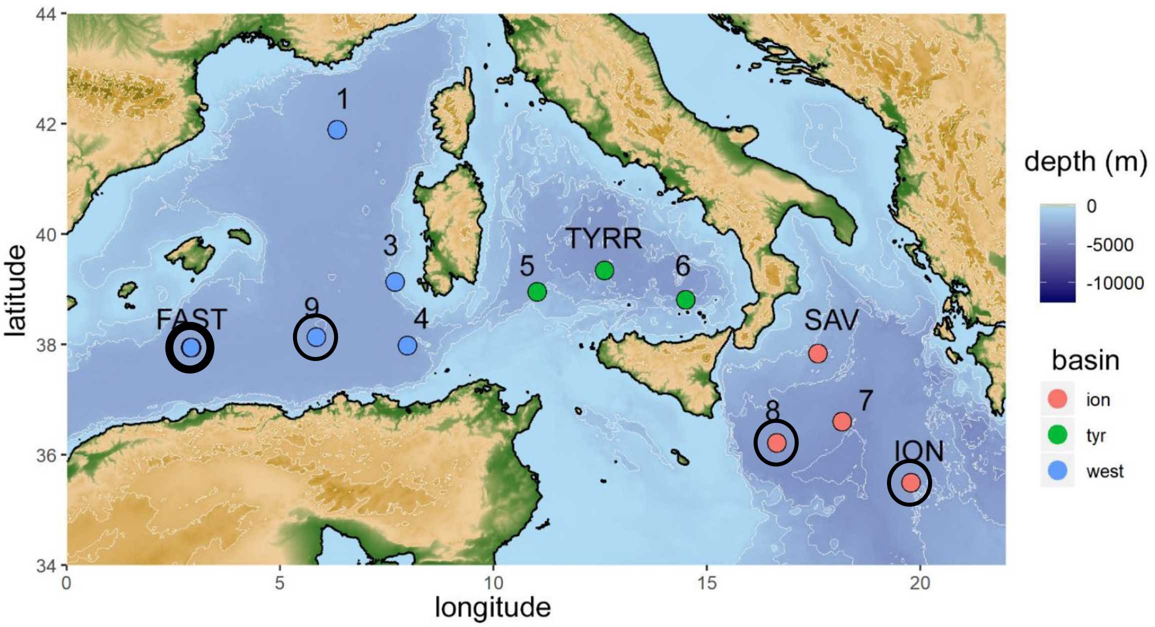


**Figure S1:** Map of the stations sampled for this study during the Peacetime cruise (Modified from Zäncker *et al.,* 2021). All stations were sampled for sea surface microlayer and surface water bacterial community composition. The stations where sea spray aerosols were produced, and bacterial community composition sequenced are circled in Black. The long stations TYRR, ION and FAST were visited for several days. At TYRR and ION, samples for surface water were taken at day 1 and 2. At station FAST we sampled a in situ wet dust deposition event, surface water was sampled before the rain, 2h, 24h and 48h after the rain, and nascent sea spray bacterial community was sequenced 2h and 24h after the rain.

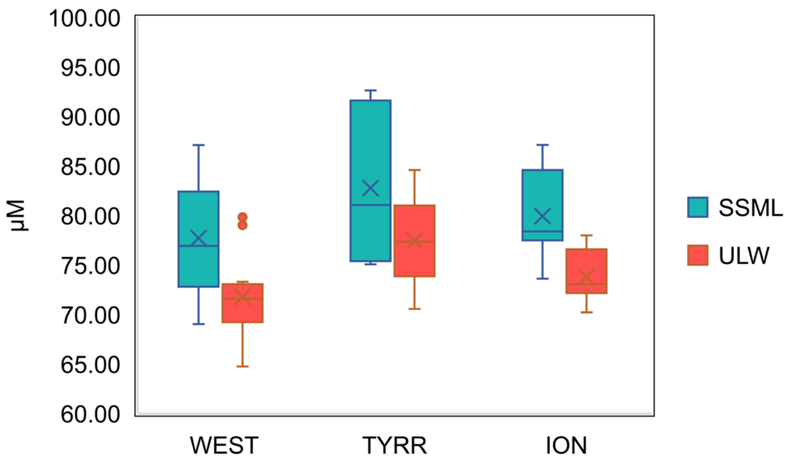


**Figure S2.** DOC concentration in the SSML and ULW in the three studied Basins.

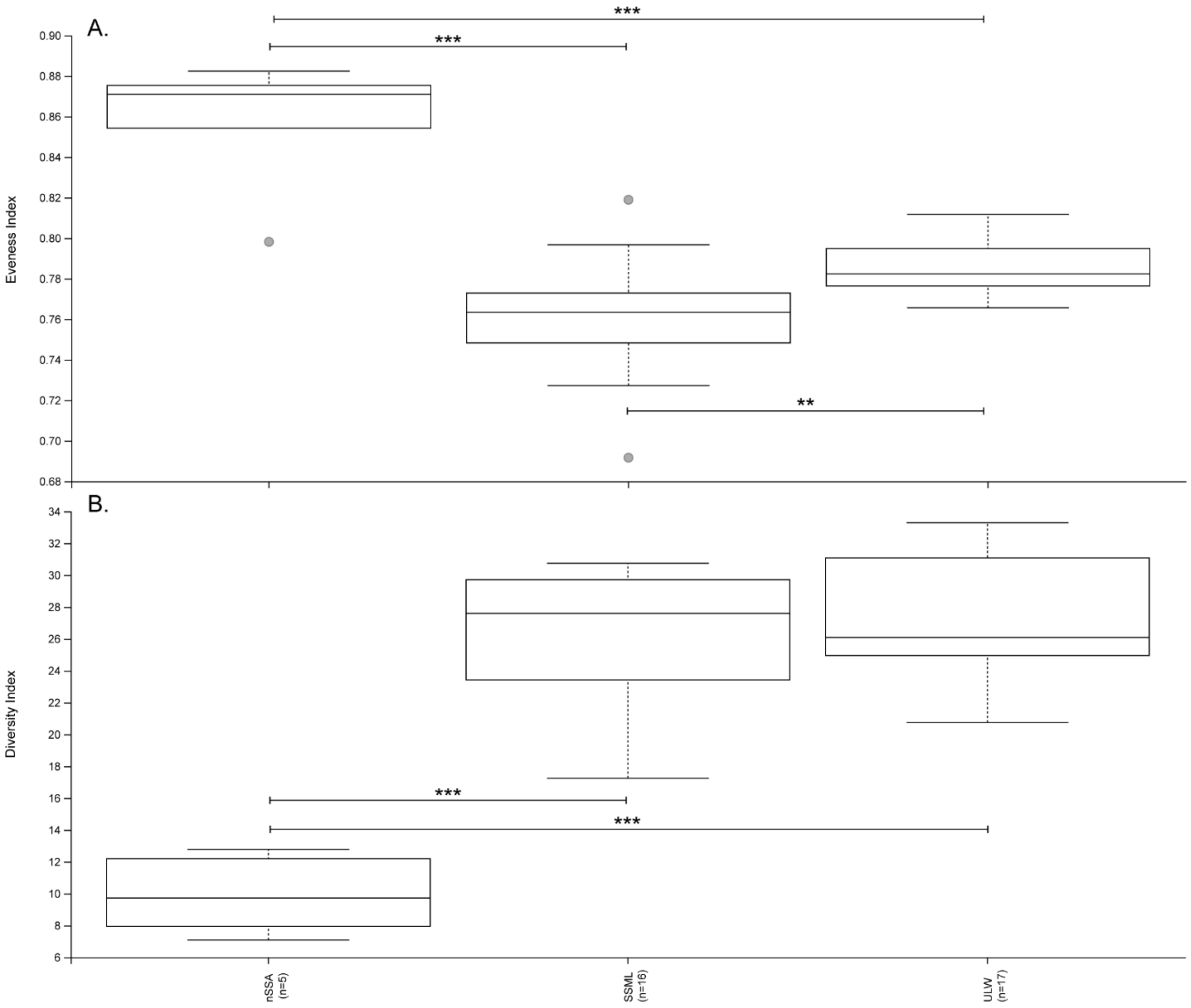


**Figure S3:** Eveness (Pielou Index, A) and Phylogenetic diversity (Faith Index, B) in the different fractions. Significant differences are shown between samples through pairwise comparison by Kurskal-Wallis test ***p <0.001,  **p=0.006. Diversity between SSML and ULW was not significantly different.

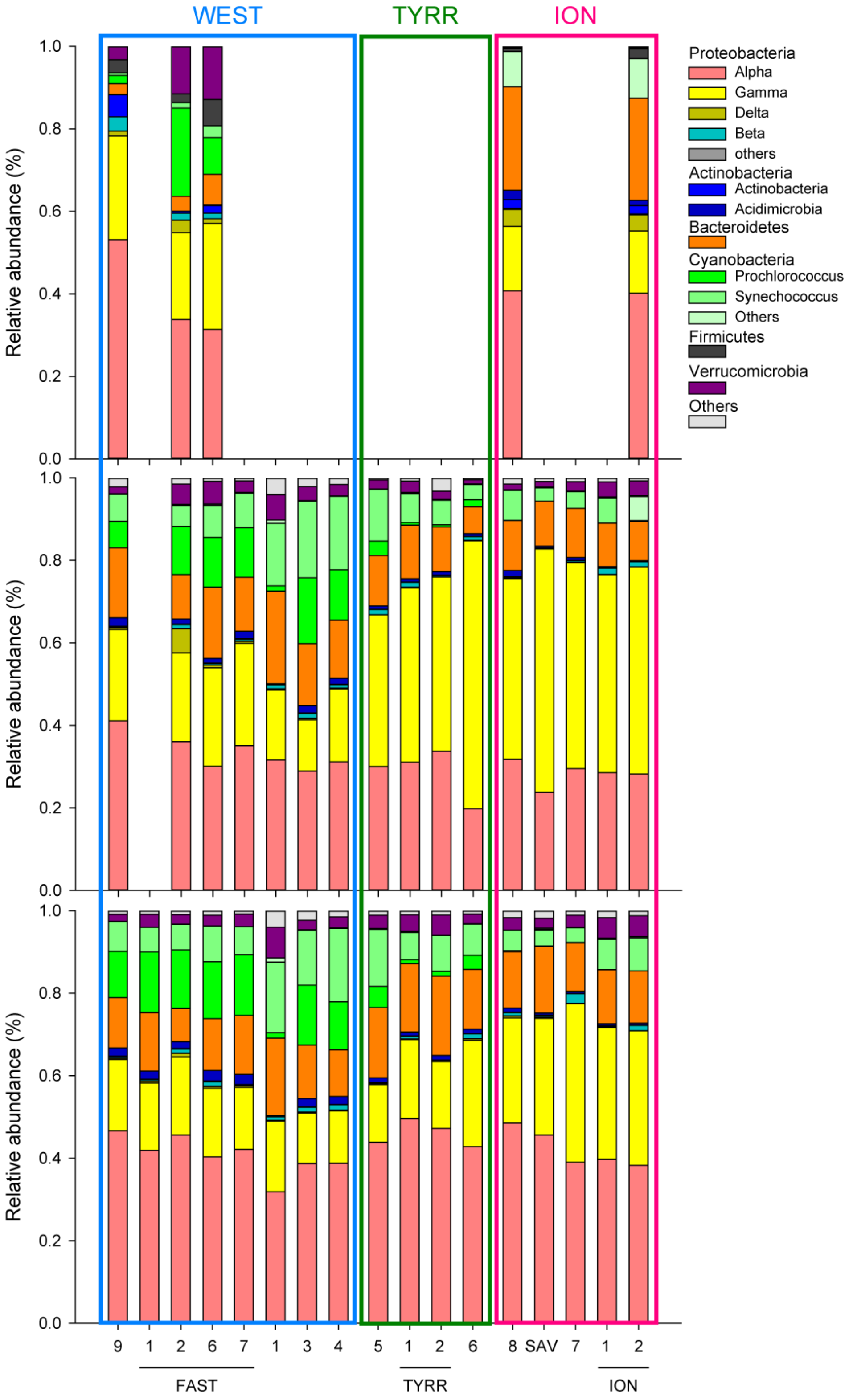


**Figure S4:** Community composition at the phylum level at each station sequenced, in the nSSA (top), SSML (middle) and ULW (bottom)

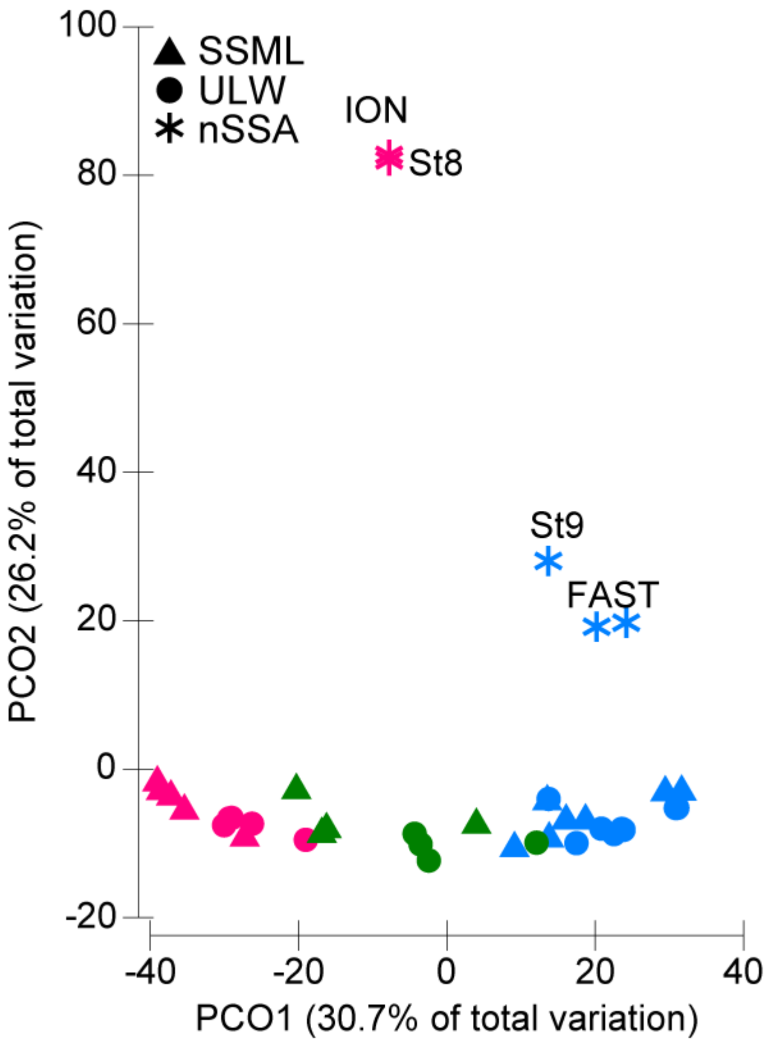


**Figure S5**: Principal component analysis of the bacterial community composition in the three different fractions: nSSA, SSML and ULW at each sequenced station based on Bray-Curtis dissimilarities of the 16S rDNA sequences. Stations are color coded per basins (Blue: Western Basin, Green: Tyrrhenian Sea and Pink: Ionian Sea)
